# Negative catalysis by the editing domain of class I aminoacyl-tRNA synthetases

**DOI:** 10.1101/2021.10.18.464777

**Authors:** Igor Zivkovic, Kate Ivkovic, Nevena Cvetesic, Ita Gruic-Sovulj

## Abstract

Aminoacyl-tRNA synthetases (AARS) translate the genetic code by loading tRNAs with the cognate amino acids. The errors in amino acid recognition are cleared at the AARS editing domain through hydrolysis of misaminoacyl-tRNAs. This ensures faithful protein synthesis and cellular fitness. Using *Escherichia coli* isoleucyl-tRNA synthetase (IleRS) as a model enzyme, we demonstrated that the class I editing domain clears the non-cognate amino acids well-discriminated at the synthetic site with the same rates as the weakly-discriminated fidelity threats. This unveiled low selectivity suggests that evolutionary pressure to optimize the rates against the amino acids that jeopardize translational fidelity did not shape the editing site. Instead, we propose that editing was shaped to safeguard cognate aminoacyl-tRNAs against hydrolysis. Misediting is prevented by the residues that promote negative catalysis through destabilisation of the transition state comprising cognate amino acid. Such powerful design allows broad substrate acceptance of the editing domain along with its exquisite specificity in the cognate aminoacyl-tRNA rejection. Editing proceeds by direct substrate delivery to the editing domain (*in cis* pathway). However, we found that class I IleRS also releases misaminoacyl-tRNA^Ile^ and edits it *in trans*. This minor editing pathway was up to now recognized only for class II AARSs.

## INTRODUCTION

Aminoacyl-tRNA synthetases (AARS) couple cognate amino acid and tRNA pairs for protein biosynthesis. They are divided into two, evolutionary distinct classes, class I and class II (1, 2). In both classes, the pairing occurs at the synthetic active site by the same two-step mechanism bearing some class-dependent features (3). The first step, amino acid activation, comprises the formation of aminoacyl-AMP (AA-AMP) while the second step is the transfer of the aminoacyl moiety to the tRNA (formation of aminoacyl-tRNA, AA-tRNA) **(Figure 1, paths 1 and 4)**. The coupling of non-cognate substrates leads to mistranslation, which can be toxic for the cell (4–6). Due to physicochemical similarities of cellular amino acids, around half of AARSs cannot achieve the tolerable level of fidelity (estimated to be 1 in 3300 (7)) in the synthetic reactions alone and thus have evolved editing (reviewed in (8, 9)). The error can be corrected by hydrolysis of non-cognate AA-AMP within the confines of the synthetic site (pre-transfer editing, **Figure 1, paths 2 and 3**) (10, 11) and/or by hydrolysis of misaminoacyl-tRNA at the dedicated editing domain (post-transfer editing) (12, 13). The latter appears to be the dominant pathway, operating by two possible routes – *in cis* **(Figure 1, path 5 and 6)** and *in trans* **(Figure 1, path 7 - 9)** (9). Editing *in trans,* so far demonstrated only in class II AARS (14), entails dissociation of the AA-tRNA and its rebinding with the 3’-end facing the editing domain.

**Figure 1.**
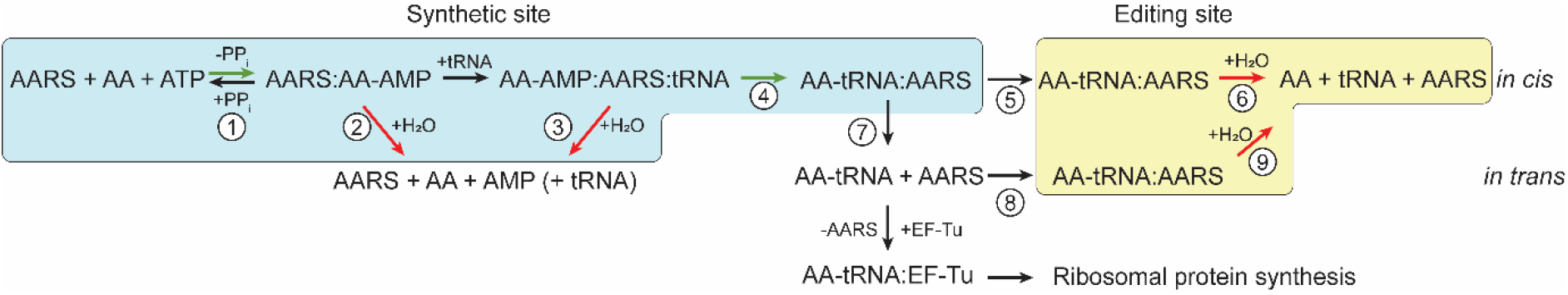
IleRS pathways of aminoacylation (green arrows) and editing (red arrows). The synthetic pathway consists of amino acid activation (1) and the aminoacyl transfer step (4). The editing pathways include tRNA-independent (2) and tRNA-dependent (3) pre-transfer editing and post-transfer editing (6, 9). Post-transfer editing can occur by translocation of AA-tRNA (5) to the editing domain for hydrolysis (6, *in cis*) or by AA-tRNA dissociation (7), its subsequent rebinding to the editing site (8) and hydrolysis (9, *in trans*).

The interplay between the synthetic and editing sites was firstly addressed by Fersht’s double-sieve hypothesis proposed originally for class I isoleucyl-(IleRS) and valyl-tRNA synthetases (ValRS) (15). It states that the synthetic site uses steric clash to discard larger than the cognate amino acids while the editing site clears smaller/isosteric non-cognate amino acids that were successfully aminoacylated to the tRNA. The steric clash was also proposed to prevent the binding of the cognate AA-tRNA to the editing domain. But, does the productive recognition at the editing site correlate well with the amino acid misrecognition at the synthetic site, and to what extent does the steric clash define the selectivity of the editing site? The former was anticipated but not experimentally addressed. The latter was tested to show that the selectivity against the cognate AA-tRNA arises from its imposed unproductive binding (16–18).

IleRS rapidly hydrolyses tRNA^Ile^s misaminoacylated with non-proteinogenic norvaline (Nva) and Val (6). This is expected as both Nva and Val are misactivated with a frequency that is 10-fold higher than the estimated tolerable error (7) and thus pose threats to the fidelity of Ile-tRNA^Ile^ formation (6, 15). Surprisingly, IleRS can also efficiently hydrolyse tRNA^Ile^ misaminoacylated with a non-proteinogenic α-aminobutyrate (Abu) and its synthetic γ-fluorinated analogues (_2_Abu and F_3_Abu), which are misactivated with up to a 20-fold *lower* frequency than the estimated tolerable error (19). This questions whether the editing site substrates need to be well misrecognized at the synthetic site, as anticipated.

Here, we set out to explore what shaped the selectivity of class I editing site and to unravel whether the same mechanisms and demands for selectivity are shared between the editing and the synthetic sites using IleRS as a model enzyme. We characterized amino acid activation and AA-tRNA^Ile^ hydrolysis using a range of amino acids with different physicochemical properties (Ala, Ser, Thr, Met, Leu, Nle). We found that IleRS synthetic site discriminates with at least 20 000-fold against the tested non-cognate amino acids. Thus, these substrates should not pose a fidelity problem. Nevertheless, all misaminoacylated tRNA^Ile^s were rapidly hydrolysed (35-65 s^−1^) at the editing site. Only cognate Ile-tRNA^Ile^ was weakly hydrolysed, demonstrating that evolution of the editing site was driven by negative catalysis (20, 21) *i.e.* selection towards destabilisation of the transition state for the cognate AA-tRNA hydrolysis (misediting). We also found that negative determinants for misediting vary among the closely related class I editing domains. Finally, we discovered that in IleRS, delivery of the AA-tRNA to the editing domain entails the accumulation of free AA-tRNA in solution, reminiscent of class II AARSs editing *in trans*.

## MATERIAL AND METHODS

### Purification of IleRS, LeuRS, and ValRS

IleRS (EC 6.1.1.5) variants were produced using QuikChange (Agilent) mutagenesis and mutations were confirmed by sequencing. Genes for *E*. *coli* IleRS (wild-type and mutants), LeuRS (EC 6.1.1.4) and ValRS (EC 6.1.1.9) inserted into pET28b plasmid were overexpressed in *E*. *coli* BL21(DE3) and purified by affinity chromatography on Ni-NTA resin (Cytiva) as described (11, 22). IleRS and ValRS were additionally purified to remove AA-AMP which is copurified bound in the enzyme active site as described (19).

### Purification and activation of the EF-Tu

The elongation factor Tu (EF-Tu, EC 3.6.5.3) with the C-terminal His-tag was overexpressed in *E*. *coli* BL21(DE3). Cells were grown to OD_600_ of 0.6-0.8 at 37 °C and expression was induced with 0.2 mM IPTG for 3 h. EF-Tu was purified on Ni-NTA resin (Cytiva) as described previously (23). EF-Tu was stored as the inactive GDP-bound form at −20 °C in a buffer containing 50 mM Hepes pH = 7.5, 10 mM MgCl_2_, 50 mM KCl, 50 % glycerol, 50 μM GDP and 5 mM β-mercaptoethanol. Activation of the EF-Tu:GDP was performed in 70 mM Hepes pH = 7.5, 50 mM ammonium acetate, 10 mM magnesium acetate, 30 mM KCl, 0.8 mM DTT, 10 mM phosphoenolpyruvate, 1 mM GTP and 0.08 U/μL pyruvate kinase (Sigma) at 37 °C for 2 hours. EF-Tu:GTP was used immediately after the activation. The activation is not efficient and results in about 10-15 % of the total EF-Tu being capable of AA-tRNA binding (23–25). Thus, the herein reported concentrations of EF-Tu presents the form capable of binding AA-tRNA (10-15 % total concentration of EF-Tu).

### Purification and labelling of tRNAs

Synthetic genes for tRNA^Ile^_GAT_ (with G1–C72 instead of WT A1–U72 sequence), tRNA^Leu^_TAA_ and tRNA^Val^_TAC_ inserted into the pET3a plasmid were overexpressed in *E*. *coli* BL21(DE3) (11, 22). Cells were grown to OD_600_ of 0.5-0.6 at 37 °C and expression was induced with 1 mM IPTG overnight at 30 °C. Substitution of the first base pair enhances transcription and does not affect tRNA^Ile^ participation in the IleRS synthetic and editing reactions (26, 27). tRNAs were isolated and purified by phenol/chloroform extraction, PEG_8000_ precipitation (removal of high molecular weight nucleic acids) and ethanol precipitation as described previously (11). Purified tRNA^Leu^ and tRNA^Val^ had acceptor activity > 90 %. The acceptor activity of tRNA^Ile^ was around 50 % so it was subjected to further purification by reverse phase chromatography on a semi-preparative Jupiter C4 column (Phenomenex), as described (11), which increased the acceptor activity to 80-90 %. tRNAs were stored in 5 mM Hepes pH 7.5. Before further use tRNA^Leu^ and tRNA^Val^ were renaturated by heating at 85 °C for 3 min, adding an equal volume of pre-heated 20 mM MgCl_2_, and slow cooling to room temperature for about 1 h. Labelled [^32^P]tRNAs were prepared as described (28, 29). [^32^P]tRNAs had acceptor activity 75-90%.

### Preparation of misaminoacylated tRNAs

Misaminoacylated [^32^P]tRNAs were prepared by mixing 25 μM tRNA^Ile^ with 5 μM T243R/D342A IleRS (mutant inactive in post-transfer editing) and a particular amino acid at the following concentration (4 mM Ala or Met; 2 mM Val, Nle or Thr; 0.2 mM Leu; 10 mM Ser) in a buffer containing 20 mM Hepes pH 7.5, 10 mM MgCl_2_, 150 mM NH_4_Cl, 2 mM ATP, 0.008 U/μL thermostable inorganic pyrophosphatase (TIPP, Sigma), 0.01 mg/mL BSA (New England Biolabs). All amino acids were purchased from Sigma. Amino acids were added in a moderate amount to prevent aminoacylation with possible Ile contaminations in the non-cognate amino acid samples. Reactions were quenched after 30 min at 37 °C by mixing with an equal amount of phenol/chloroform. AA-tRNA^Ile^s were purified by phenol/chloroform extraction followed by two consecutive steps on Bio-Spin P30 columns (Bio-Rad) and dialyzed against 10 mM NaOAc pH 4.5. Before further use AA-tRNA^Ile^s were renaturated by heating at 85 °C for 3 min, adding an equal volume of pre-heated 20 mM MgCl_2_, and slow cooling (around 1 h) to room temperature.

### Amino acid activation

Amino acid activation was followed by an ATP-PP_i_ exchange assay (30–32) which was performed at 37 °C in a buffer containing 50 mM Hepes pH 7.5, 20 mM MgCl_2_, 0.1 mg/mL BSA, 5 mM DTT, 4 mM ATP and 1 mM [32P]PP_i_ (Perkin-Elmer). Enzymes were present at 50 to 100 nM while amino acid concentrations were varied from 0.1 to 10 × *K*_M_. Reactions were quenched by mixing 1.5 μL of the reaction mixture with 3 μL of the quench solution (600 mM NaOAc pH 4.5 and 0.15 % SDS). Formed [^32^P]ATP was separated from the remaining [^32^P]PP_i_ by thin-layer chromatography (TLC) on polyethyleneimine plates (Macherey-Nagel) in 4 M urea and 750 mM KH_2_PO_4_ pH 3.5. Signal visualization was performed on a Typhoon Phosphoimager (GE Healthcare) and quantified with ImageQuant software as described (29). Kinetic parameters (*k*_cat_ and *k*_sp_) were obtained by fitting the data to the modified Michalis-Menten equation: 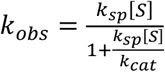, where *k*_cat_ is the turnover number, *k*_sp_ is the specificity constant (*k*_cat_/*K*_M_), [S] is the amino acid concentration and *k*_obs_ is the observed rate constant. *k*_obs_ is calculated as *v*_0_/[E]_0_, where *v*_0_ is the initial reaction rate and [E]_0_ is the enzyme concentration. The data were fitted using GraphPad Prism software. We opted for the modified Michaelis-Menten equation due to the benefits discussed recently by K. Johnson (33). Briefly, by fitting the data to the modified model, *k*_sp_ is obtained directly, rather than being calculated from *k*_cat_ and *K*_M_, which leads to smaller errors.

### Two-step aminoacylation

Aminoacylation of tRNA^Ile^, comprising both activation and the transfer step, by various IleRS variants was followed at 37 °C in a buffer containing 20 mM Hepes pH 7.5, 10 mM MgCl_2_, 150 mM NH_4_Cl, 2 mM ATP, 0.008 U/μL TIPP and 0.01 mg/mL BSA, 15 μM tRNA^Ile^ 1 mM Ile and 20 nM WT, T246A, H333A, H333G, T246A/H333A or L247A/H333G IleRS. Misaminoacylation of tRNA^Ile^ was followed under the same conditions except that the concentrations of enzymes were higher (1 μM WT or T243R/D342A IleRS) and non-cognate amino acids were used at following concentrations: Leu 0.2 mM; Nle and Thr 2 mM; Ala and Met 4 mM; Ser 10 mM. The reactions were stopped by mixing 1.5 μL of the reaction mixture with 3 μL of the quench solution (600 mM NaOAc pH 4.5 and 0.15 % SDS). 1.5 μL of the quenched reaction mixture was mixed with 3 μL of P1 nuclease (Sigma) (≥0.01 U/μL in 300 mM NaOAc pH 5.0 and 0.15 mM ZnCl_2_) and incubated for 1 hour at room temperature. The P1 nuclease treatment releases terminal adenine nucleotide, with or without amino acid attached as the result of tRNA aminoacylation. Free and aminoacylated [^32^P]AMP were separated by TLC in 100 mM NaOAc and 5 % HOAc. Signal visualization was performed on a Typhoon Phosphoimager (GE Healthcare) and quantified with ImageQuant software as described (29).

### Parallel Formation of AMP and AA-tRNA

Formation of [^32^P]AMP and AA-[^32^P]tRNA were followed at 37 °C in parallel reactions, in a buffer containing 50 mM Hepes 7.5, 20 mM MgCl_2_, 0.1 mg/mL BSA and 2 mM DTT supplemented with 0.1 mg/mL BSA, 1 mM ATP and 0.004 U/μL thermostable inorganic pyrophosphatase and 10-12 μM tRNA. The reactions that monitored [^32^P]AMP formation were supplemented with [α-^32^P]ATP (0.01-0.1 mCi/mL) (Perkin Elmer) while the reactions that monitored AA-[^32^P]tRNA were supplemented with [^32^P]tRNA (0.01-0.1 mCi/mL). The enzymes were 2 μM and amino acids were used at the following concentrations: 2 mM Ile, 20 mM Val and 30 mM Nva or Thr. The concentration of the GTP-bound EF-Tu, when added, was estimated to 8-12 μM (total concentration of added EF-Tu was 80 μM). The reactions were stopped by mixing 1.5 μL of the reaction mixture with 3 μL of the quench solution (600 mM NaOAc pH 4.5 and 0.15 % SDS). For following AMP formation, [^32^P]ATP and [^32^P]AMP were separated by TLC in 100 mM NaOAc and 5 % HOAc. When the formation of AA-tRNAs was followed, the quenched reaction mixtures were degraded by P1 nuclease and separated as described for the two-step aminoacylation (see above). Signal visualization was performed on a Typhoon Phosphoimager (GE Healthcare) and quantified with ImageQuant software as described (29).

### Single-turnover hydrolysis

Single-turnover hydrolysis of AA-tRNAs was performed at 37 °C by mixing equal volumes of 20 μM IleRS in a buffer containing 200 mM Hepes pH 7.5, 75 mM NH_4_Cl, 20 mM MgCl_2_, 5 mM DTT and 0.01 mg/mL BSA and freshly renaturated misaminoacylated [^32^P]tRNA^Ile^ (0.2-1 μM) in 10 mM NaOAc pH 4.5 as described previously (19). Mixing of the enzyme and AA-[^32^P]tRNAs was done using a rapid chemical quench instrument (RQF-3, KinTek Corp.). For the reactions having *t*_1/2_ ≥ 5 s, manual mixing was performed. The reaction mixtures were quenched, treated with P1 nuclease, AA-[^32^P]AMP and [^32^P]AMP were separated and analysed as described for the aminoacylation (see above and (29)). The data were fitted to the single exponential equation 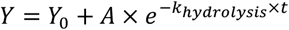, where Y_0_ is the y intercept, A is a scaling constant, *k*_hydrolysis_ is the apparent hydrolytic rate constant, and *t* is time.

### Effect of the expression of IleRS variants on the growth of *E. coli*

*E. coli* BL21(DE3) strain was transformed with pET28 plasmids carrying genes for either WT IleRS, H333A IleRS or T246A/H333A IleRS. The empty plasmid was used as a control. The cultures were grown in 100 mL culture flasks in the M9 media with the addition of 0.4 % glucose and 30 μg/mL kanamycin. Overnight cultures were diluted to OD_600_ of 0.04 and supplemented with 100 μM IPTG to induce the protein expression. The growth at 37 °C and 250 rpm was monitored using UV-Vis spectrophotometer Evolution 60S (Thermo Scientific). The expression profile was followed by SDS-PAGE. The data were fitted to the logistic model 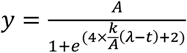, where *A* is the maximal cell growth, *λ* is the lag phase and *k* is the growth rate constant.

## RESULTS

### The editing site clears a broad range of misaminoacylated tRNA^Ile^s

Misaminoacylation of tRNA^Ile^ with the non-cognate amino acids that are significantly different from Ile (**Supplementary Figure S1**) is challenging. However, this is a prerequisite for addressing the selectivity of the editing site as post-transfer editing is tested by following hydrolysis of preformed misaminoacylated tRNAs (29). In doing so, we found that all tested non-cognate amino acids were activated (**Figure 1, path 1**) albeit with high discrimination factors (*D* > 20 000, **Table 1**), which reflect efficient exclusion of the non-cognate substrate from the IleRS synthetic site. This is in accordance with known high selectivity of the IleRS synthetic site (6, 19, 34–38). Somewhat surprisingly, despite weak misactivation, tRNA^Ile^ was successfully misaminoacylated (up to 60 % aminoacylation level) with all tested non-cognate amino acids by post-transfer editing deficient T243R/D342A IleRS (6, 11, 19) (**Supplementary Figure S2** and **Supplementary Figure S3**). Interestingly, we did not observe aminoacylation with WT IleRS (**Supplementary Figure S2, inset**). Thus, the editing deficient AARSs can provide an alternative to the ribozyme approach (39) for tRNA misaminoacylation. This analysis is further complicated by artefacts that may arise from contamination of non-cognate amino acid samples with trace amounts of the cognate amino acids (5, 15, 40). For that reason, we estimated the purity of the used amino acids (**Supplementary Figure S4**) and found that Leu, and possibly also Met and Nle, may contain trace amounts of Ile.

**Table 1.** Kinetic parameters for activation of amino acids by WT IleRS.

| Amino acid | $k_{cat} / s^{-1}$ | $k_{sp}^a / s^{-1} mM^{-1}$ | $K_M / mM$ | $D^b$ |
| --- | --- | --- | --- | --- |
| Ile | $56.7 \pm 0.3$ | $(16.6 \pm 0.4) \times 10^3$ | $(3.41 \pm 0.06) \times 10^{-3}$ | 1 |
| Val<sup>c</sup> | $36 \pm 6$ | 77 | $0.47 \pm 0.03$ | 156 |
| Nva<sup>d</sup> | $41 \pm 1$ | 50 | $0.82 \pm 0.07$ | 239 |
| Abu<sup>e</sup> | $23 \pm 3$ | $1.6 \pm 0.1$ | $15 \pm 1$ | 10 375 |
| Thr | $32 \pm 2$ | $0.82 \pm 0.04$ | $39 \pm 2$ | 20 243 |
| F<sub>2</sub>Abu<sup>e</sup> | $6.9 \pm 0.7$ | $0.45 \pm 0.8$ | $16 \pm 2$ | 36 888 |
| F<sub>3</sub>Abu<sup>e</sup> | $5.3 \pm 0.8$ | $0.19 \pm 0.02$ | $28 \pm 2$ | 87 368 |
| Ala | $10 \pm 1$ | $0.10 \pm 0.02$ | $100 \pm 9$ | 166 000 |
| Ser | - | $0.016 \pm 0.005^f$ | - | 1 037 500 |
| Met | $10.6 \pm 0.4$ | $7.6 \pm 0.7$ | $1.4 \pm 0.1$ | NC<sup>g</sup> |
| Nle | $4.8 \pm 0.4$ | $1.6 \pm 0.2$ | $3.1 \pm 0.6$ | NC<sup>g</sup> |
| Leu | $28.7 \pm 0.7$ | $26 \pm 3$ | $1.1 \pm 0.1$ | NC<sup>g</sup> |

The activation step was tested by ATP-PPi exchange assay. The values represent the average value ± SEM of at least three independent experiments.
<sup>a</sup> $k_{sp}$ – specificity constant ( $k_{cat}/K_M$ ) is obtained from the modified Michaelis-Menten equation $k_{obs} = \frac{k_{sp}[S]}{1 + \frac{k_{sp}[S]}{k_{cat}}}$ (33).
<sup>b</sup>Discrimination factor – $k_{sp,cognate}/k_{sp,non-cognate}$ , *i.e.* $(k_{cat}/K_M)_{cognate}/(k_{cat}/K_M)_{non-cognate}$ . The values below 3300 indicate higher error frequencies than the estimated error of protein synthesis (7).
<sup>c</sup>Data was taken from (11). The $k_{cat}$ and $K_M$ values were determined using the unmodified form of the Michaelis-Menten equation. $k_{sp}$ was calculated by dividing $k_{cat}$ with $K_M$ .
<sup>d</sup>Data was taken from (6). The $k_{cat}$ and $K_M$ values were determined using the unmodified form of the Michaelis-Menten equation. $k_{sp}$ was calculated by dividing $k_{cat}$ with $K_M$ .
<sup>e</sup>Raw data was taken from (19).
<sup>f</sup> $k_{cat}$ and $K_M$ were not determined due to the low activity.
<sup>g</sup>Not calculated due to possible contamination of the amino acid sample with cognate Ile.

Next, we isolated the post-transfer editing step by mixing preformed misaminoacylated tRNA^Ile^ with a surplus of WT IleRS, using a rapid chemical quench instrument. The hydrolysis of misaminoacylated tRNA^Ile^ was followed in time to calculate the first-order rate constant (**Supplementary Figure S3**). The single-turnover conditions ensure that product dissociation does not limit the observed rate (22). A 2-fold higher concentration of IleRS or AA-tRNA^Ile^ returned the same hydrolysis rate confirming that binding is not rate-limiting. Thus, the observed rate constants (**Figure 2**, **Supplementary Figure S3**) represent the catalytic step (hydrolysis of misaminocylated tRNA^Ile^) within the editing site.

**Figure 2.**
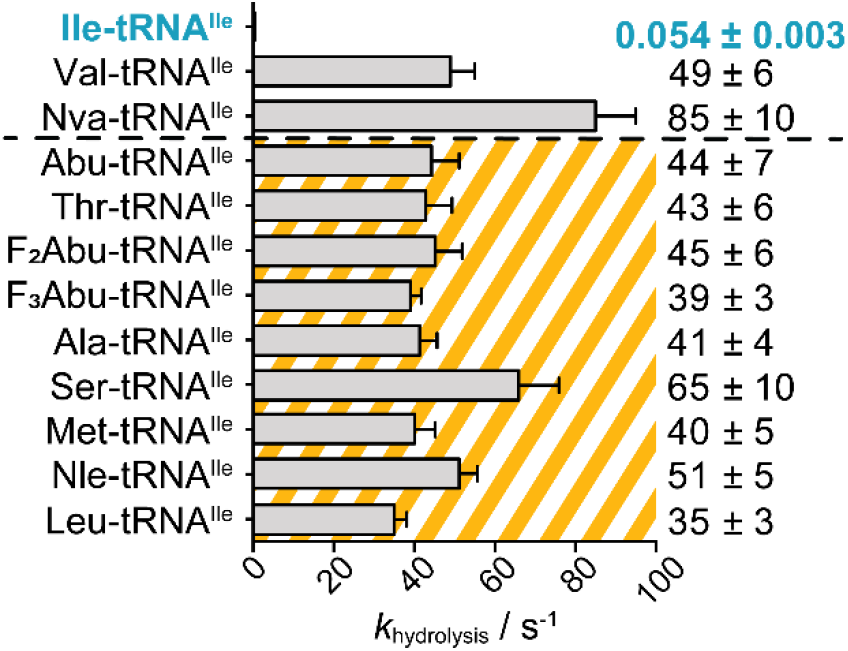
Single-turnover hydrolysis of misaminoacylated tRNAs by WT IleRS. tRNAs misaminoacylated with amino acids that are well discriminated at the IleRS synthetic site (*D* > 3300) are presented in the striped area. Value 3300 is taken as the tolerable error of protein synthesis was estimated around 1 in 3300 (7). Rapid hydrolysis of Leu-, Met-, and Nle-tRNA^Ile^ confirmed that possible traces of cognate Ile in the Leu, Met or Nle samples did not compromise the editing analysis. Time courses from which the first-order rate constants (*k*_hydrolysis_) were calculated are presented in Supplementary Figure S3. *k*_hydroysis_ for Val-, Nva-, Abu-, F_2_Abu-, and F_3_Abu-tRNAsIle were taken from (6, 19).

The single-turnover analysis revealed, in agreement with the incapacity of the WT IleRS to accumulate misaminoacylated tRNAs^Ile^ (**Supplementary Figure S2, inset**), that all misaminoacylated-tRNA^Ile^s were rapidly hydrolysed with similar rates ranging from 35 to 65 s^−1^ (**Figure 2**, please note that a possible Leu, Met and Nle contaminations did not compromise the editing analysis). This is surprising as these amino acids (except Val and Nva) are efficiently discriminated at the synthetic site and as such cannot pose a threat to IleRS aminoacylation fidelity. Finding that amino acids are rapidly cleared at the editing domain irrespectively of the requirement for their editing, lend a new paradigm about the editing selectivity principles. Moreover, the editing site shows no clear preference towards physicochemical features of the editing substrates, like size (Met and Nle, both with longer unbranched side chain, are eliminated), hydrophobicity (polar Ser and Thr are efficiently cleared at the editing site) or branching (Leu-tRNA^Ile^ is also rapidly hydrolysed). Cognate Ile-tRNA^Ile^ was the only exemption, suggesting, that prevention of cognate AA-tRNA misediting was a major constraint during the evolution of the editing site. Thus, we set to explore how the editing site excludes the cognate Ile-tRNA^Ile^ and in parallel promotes editing of misaminoacylated tRNAs.

### Negative determinants for Ile-tRNA^Ile^ misediting

Two conserved residues of the editing domain, Thr246 and His333 (Thr233 and His319 in *T. thermophilus* IleRS, PDB ID: 1WNZ, **Supplementary Figure S5**) were previously characterised by time-course analysis and their equal contribution to the rejection of Ile-tRNA^Ile^ has been proposed (41, 42). However, in closely related LeuRS, the specificity against the cognate Leu resides solely on Thr252 (analogous to Thr246 in EcIleRS) (16). It has been shown that time-course analysis may lead to incorrect models of enzyme mechanisms (11). Therefore, we used single-turnover catalysis to assign the individual contributions of Thr246 and His333 (**Table 2)**. Interestingly, T246A substitution, increased the rate of Ile-tRNA^Ile^ misediting by only 2-fold (0.126 ± 0.006 s^−1^). In contrast, the H333A mutant showed a 20-fold increase in the rate of Ile-tRNA^Ile^ hydrolysis (1.04 ± 0.06 s^−1^), arguing that, for IleRS, His333 is the main negative determinant of misediting. Finally, the T246A/H333A mutant showed a 260-fold increase in the rate of Ile-tRNA^Ile^ hydrolysis (14 ± 1 s^−1^), displaying about 7-fold higher effect than the cumulative effects of the independent mutations. His333 was further mutated to Gly, which additionally increased the rate of Ile-tRNA^Ile^ hydrolysis (4-fold compared to H333A), suggesting that steric hindrance may contribute to the His333 action. Importantly, the mutants displayed only 2-fold slower rates of editing of non-cognate Val-tRNA^Ile^ relative to the WT (**Table 2**) pointing towards their almost exclusive effect on Ile-tRNA^Ile^ misediting. Thus, the main negative determinant of the IleRS editing site appears to be His333 whose role is synergistically supported by Thr246. This contrasts LeuRS which utilizes Thr252 as a sole negative determinant and suggests idiosyncratic evolution of the mechanisms governing rejection of the cognate product in class I editing domains.

**Table 2.** Single-turnover and steady-state rate constants for different variants of IleRS.

| Enzyme | $k_{\text{hydrolysis}}^{\text{a}} / \text{s}^{-1}$ | | $k_{\text{aminoacylation}}^{\text{b}} / \text{s}^{-1}$ |
| --- | --- | --- | --- |
|  | Ile-tRNA<sup>Ile</sup> | Val-tRNA<sup>Ile</sup> | Ile |
| WT | 0.054 ± 0.003 | 49 ± 6<sup>c</sup> | 1.3 ± 0.2 |
| T246A | 0.126 ± 0.006 | 32 ± 2 | 1.5 ± 0.3 |
| H333A | 1.04 ± 0.06 | 19.2 ± 0.6 | 1.6 ± 0.4 |
| H333G | 4.4 ± 0.7 | 24 ± 1 | 1.1 ± 0.2 |
| T246A/H333A | 14 ± 1 | 24 ± 1 | 0.96 ± 0.06 |

The values represent the average value ± SEM of at least three independent experiments.
<sup>a</sup>Single-turnover rate constants for AA-tRNA<sup>Ile</sup> hydrolysis. Time courses from which the first-order rate constants ( $k_{\text{hydrolysis}}$ ) were calculated are presented in **Supplementary Figure S6**.
<sup>b</sup>Steady-state rate constants for Ile-tRNA<sup>Ile</sup> synthesis.
<sup>c</sup>Data was taken from (6).

### IleRS deprived of the negative determinants misedits Ile-tRNA^Ile^ *in trans*

During steady-state (mis)aminoacylated AA-tRNA partitions between hydrolysis (editing; **Figure 1 path 6**) and dissociation (**Figure 1, path 7**) (product release). To reach the editing site, AA-tRNA translocates (**Figure 1, path 5**) its single-stranded 3’ end while the tRNA body remains bound to the enzyme (editing *in cis*). In case when whole AA-tRNA dissociates from the enzyme prior reaching the editing site, it can re-bind from solution with the 3’ end facing the editing domain (**Figure 1, paths 8 and 9,** editing *in trans*). Editing *in cis* depletes the product and thus compromises steady-state aminoacylation. In contrast, editing *in trans* may not affect the aminoacylation rate, because re-binding of AA-tRNA for hydrolysis is not favoured at low steady-state enzyme concentration. Therefore, the finding that both H333A and T246A/H333A IleRSs exhibit little to no change in steady-state aminoacylation rates relative to the WT enzyme (*k*_aminoacylation_, **Table 2**), despite rapid Ile-tRNAIle hydrolysis at their editing sites (*k*_hydrolysis_, **Table 2**), implies that these mutants misedit Ile-tRNAIle *in trans*. This is unexpected as editing *in trans* was not yet demonstrated for class I AARSs.

Non-stoichiometric ATP consumption is diagnostic of active editing as futile aminoacylation/editing cycles consume ATP without accumulating AA-tRNA. To address whether the IleRS mutants misedit Ile-tRNA^Ile^ *in trans*, two complementary approaches were undertaken: i) we used higher IleRS concentration (2 μM instead of 20 nM used in the steady-state aminoacylation) to favour re-binding of Ile-tRNA^Ile^ and thus misediting *in trans* and ii) higher IleRS concentration was complemented by the addition of 8-12 μM active EF-Tu, which may suppress misediting *in trans* by competing with IleRS in the binding of free Ile-tRNA^Ile^ (43).

ATP consumption (AMP formation) and Ile-tRNA^Ile^ formation were followed in parallel reactions that differ only in the labelled components – [^32^P]ATP was used for the former and [^32^P]tRNA^Ile^ for the latter (**Supplementary Figure S7**). The ratio of consumed ATP per Ile-tRNA^Ile^ accumulated in solution (AMP/Ile-tRNA^Ile^) was calculated for the reactions without and with EF-Tu (**Figure 3**). In the absence of EF-Tu, both mutants consume 18-(H333A) to 1100-fold (T246A/H333A) higher than the stoichiometric amount of ATP per released Ile-tRNA^Ile^. Thus, multi-turnover conditions at high concentrations of the mutants support misediting. That misediting takes place *in trans*, is further supported by 9-(H333A) to 18-fold (T246A/H333A) drop in AMP/Ile-tRNA^Ile^ ratio in the presence of EF-Tu. The WT enzyme, exhibiting marginal Ile-tRNA^Ile^ misediting, used stoichiometric amount of ATP per Ile-tRNA^Ile^, independently on the presence/absence of EF-Tu. Interestingly, the significant energetic cost of misediting was exhibited mainly with T246A/H333A, raising an intriguing question - how detrimental is hydrolysis of Ile-tRNA^Ile^?

**Figure 3.**
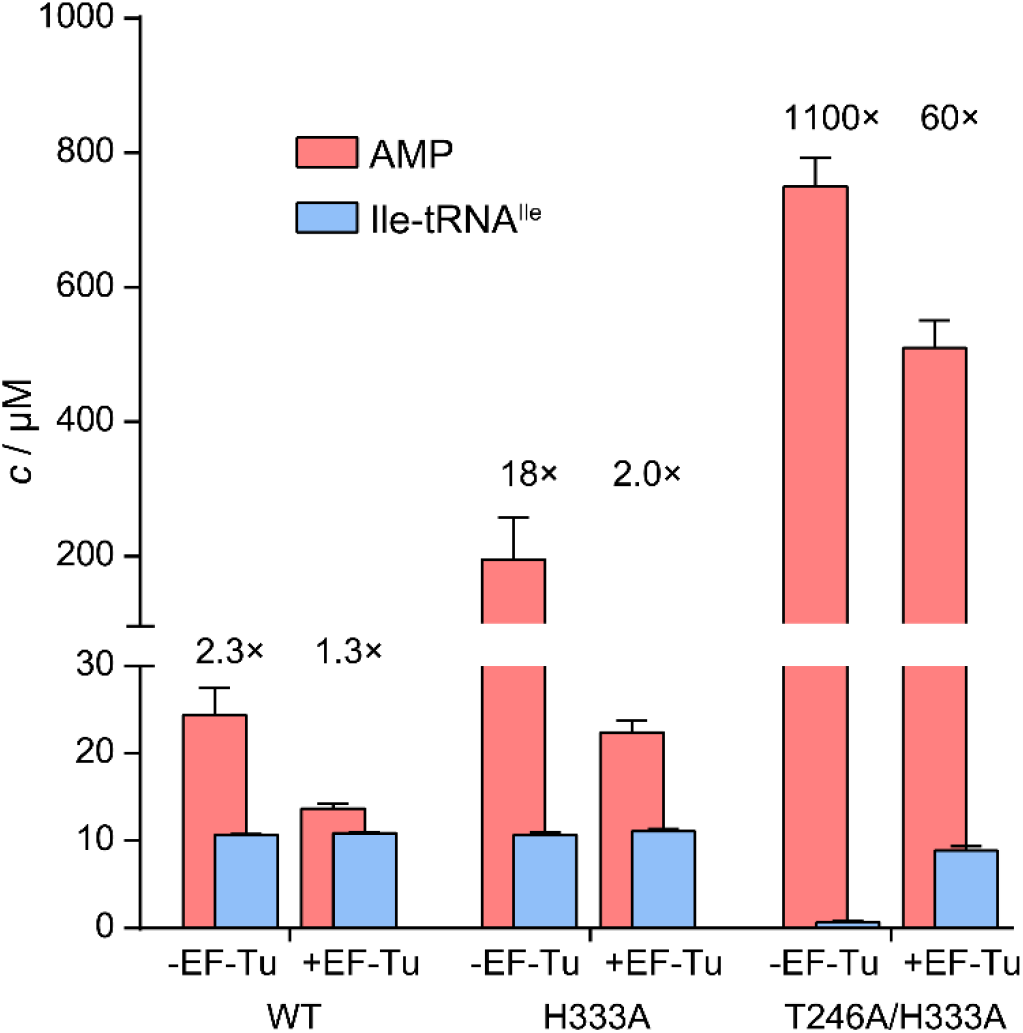
AMP (pink) and Ile-tRNA^Ile^ (blue) concentrations at the 5-minute time point for WT, H333A and T246A/H333A IleRS. The time courses are given in Supplementary Figure S7. The numbers above bars are AMP/Ile-tRNA^Ile^ ratios. The enzymes were 2 μM, the tRNA^Ile^ 12 μM, and EF-Tu (the active GTP-form) was 8-12 μM.

### Ile-tRNA^Ile^ misediting impairs cell growth

To investigate to what extent misediting of Ile-tRNA^Ile^ affects cell viability, we followed the growth of *E*. *coli* BL21(DE3) strain transformed with the plasmids encoding WT IleRS or its Ile-tRNA^Ile^ misediting active variants (H333A and T246A/H333A). A moderate expression (**Supplementary Figure S8**) of the WT enzyme did not show any growth defects demonstrating that expression *per se* is not a burden for the cell (**Figure 4**). Interestingly, the H333A mutant did not significantly influence the growth rate suggesting that Ile-tRNA^Ile^ misediting of 1 s^−1^ could be physiologically tolerated. In contrast, the T246A/H333A mutant (hydrolytic rate of 14 s^−1^) showed a noticeable growth defect (**Figure 4**), in agreement with the negative selection against this activity.

**Figure 4.**
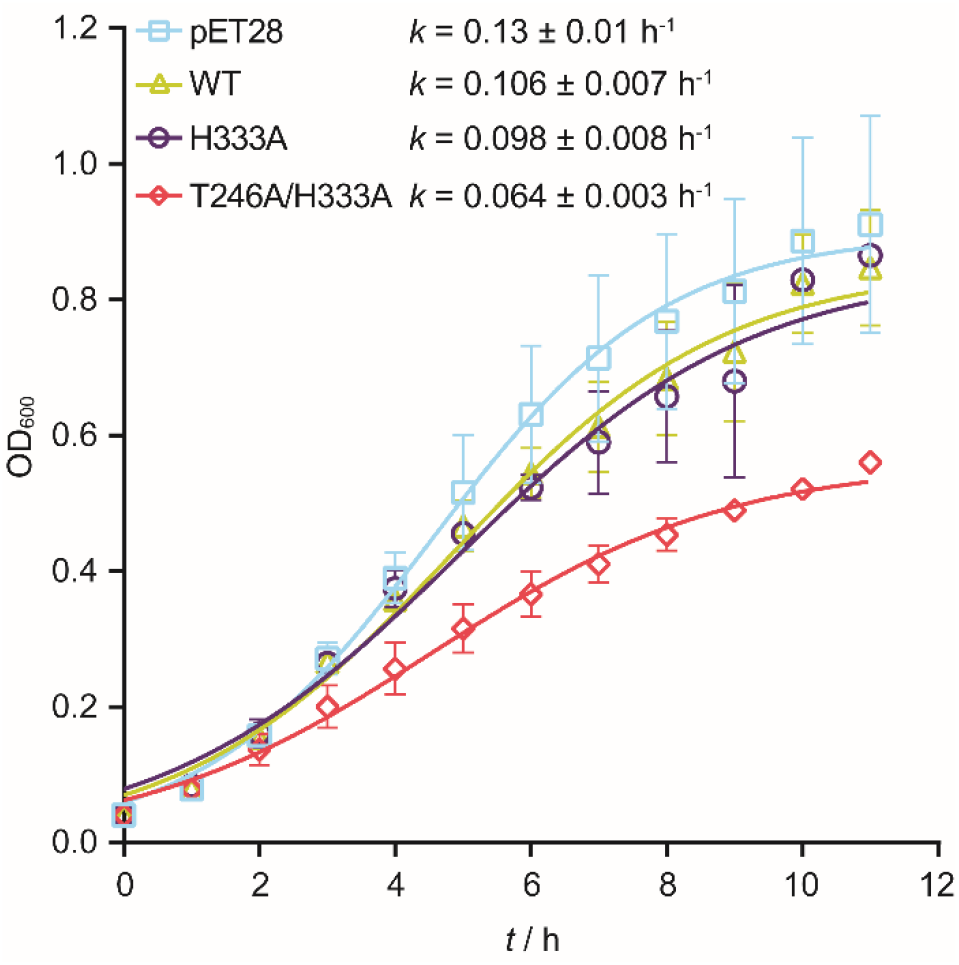
Growth of *E. coli* BL21(DE3) transformed with the pET28 plasmids carrying IleRS variants. The empty plasmid was used as a control. The growth was followed in M9 medium supplemented with 100 μM IPTG. The growth rate constant (*k*) was determined from the logistic growth model. In the absence of IPTG, the growth rate of *E. coli* BL21(DE3) transformed with the empty pET28 was 0.28 ± 0.01 h^−1^. The basal level of Ile-tRNA^Ile^ synthesis by endogenous WT IleRS should not pose a problem as Ile-tRNA^Ile^ produced by any route is subjected to editing *in trans*.

### IleRS is unique among class Ia AARSs in exhibiting a high level of editing *in trans*

To address whether IleRS edits *in trans* its biological threat Val-tRNA^Ile^, we followed the accumulation of AMP and Val-tRNA^Ile^ by the WT enzyme as described above bearing in mind that EF-Tu may bind tRNAs misaminoacylated with near-cognate amino acids (14, 23, 44). In the absence of EF-Tu, the analysis returned the AMP/Val-tRNA^Ile^ ratio of 1330 and a minor accumulation of Val-tRNA^Ile^ (**Figure 5, Supplementary Figure S9**), both in agreement with the efficient Val-tRNA^Ile^ editing (27). The addition of EF-Tu increased the accumulation of Val-tRNA^Ile^ by 10-fold and decreased the amount of consumed ATP by more than 3-fold, leading to a significant drop (58-fold) in AMP/Val-tRNA^Ile^ ratio (23 *vs* 1330). This indicates that IleRS edits Val-tRNA^Ile^ *in trans*, providing to the best of our knowledge the first demonstration of editing *in trans* for a class I AARS.

**Figure 5.**
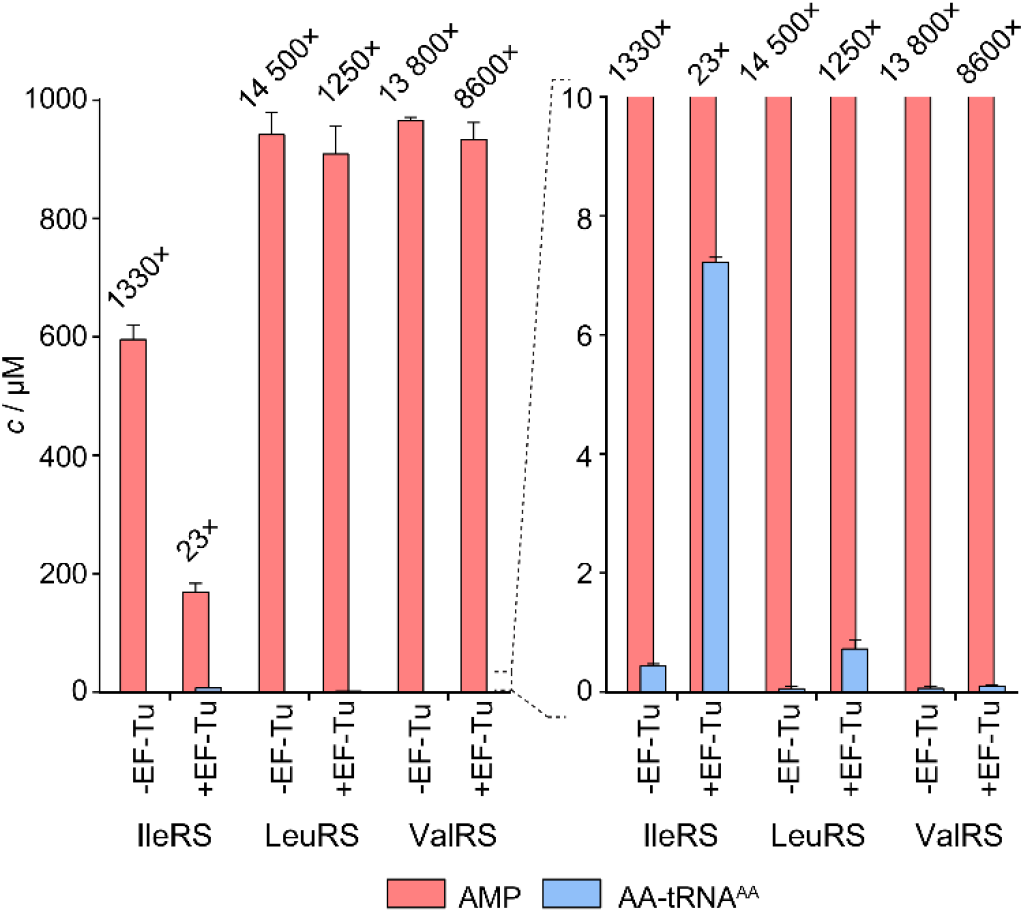
AMP (pink) and misaminoacylated tRNA (blue) concentrations at the 5-minute time point for IleRS, LeuRS and ValRS. The time courses are given in Supplementary Figure S9. The numbers above bars are AMP/Ile-tRNA^Ile^ ratios. The enzymes were 2 μM, the tRNAs 10 μM, and EF-Tu (the active GTP-form) was 8-12 μM.

Next, we tested whether LeuRS and ValRS also use editing *in trans* with their main biological threats Nva and Thr, respectively (5, 45). Both LeuRS and ValRS consumed a highly non-stoichiometric amount of ATP per accumulated misaminoacylated tRNA (14 500 and 13 800, **Figure 5, Supplementary Figure S9**), in agreement with established editing of Nva-tRNA^Leu^ and Thr-tRNA^Val^ (6, 22, 45). The addition of EF-Tu dropped the ATP/ Thr-tRNA^Val^ ratio by less than 2-fold (13 800 *vs* 8600). The lack of EF-Tu effect indicates that ValRS edits Thr-tRNA^Val^ *in cis*. The picture is more complicated for LeuRS, where the presence of EF-Tu promotes a 12-fold drop in ATP/AA-tRNA ratio (14 500 *vs* 1250) that may indicate the participation of editing *in trans*. However, the drop does not stem from a decrease in the ATP consumption, which is only 1.04-fold lower in the presence of EF-Tu. Thus, the cycles of Nva-tRNA^Leu^ hydrolysis and the subsequent tRNA^Leu^ misaminoacylation which consumes ATP are not influenced by EF-Tu. This strongly suggests that LeuRS mainly operates *in cis* in agreement with the previous data (14). The observed EF-Tu-dependent accumulation of AA-tRNA^Leu^ is puzzling and likely originates from trace contaminations of Leu in the Nva sample (**Supplementary Figure S10**). Indeed, the accumulation of Leu-tRNA^Leu^, like of Nva-tRNA^Leu^ can be diminished by rebinding to LeuRS and (mis)editing. Yet, hydrolysis of Leu-tRNA^Leu^ is 3×10^3^-times slower than Nva-tRNA^Leu^ (22), contributing minimally to the ATP consumption. EF-Tu may bind Leu-tRNA^Leu^ and thus affects its accumulation but without a noticeable effect on ATP consumption. To conclude, our data show that IleRS is distinct from closely related LeuRS and ValRS in a fraction of post-transfer editing that operates *in trans*.

## DISCUSSION

### Class I AARS synthetic and editing sites act as mirror images

AARSs are textbook examples of how high selectivity emerged under strong evolutionary pressure to evade deleterious errors (38). Their synthetic sites adopt numerous strategies to enforce recognition of the cognate and rejection of the non-cognate amino acids (46–49). If non-cognate amino acid, however, gets coupled to the tRNA, post-transfer editing resolves the problem. The editing site evolved to clear amino acids that jeopardize the accuracy of translation arguing that amino acids well discriminated at the synthetic site (large discrimination factor, *D*) will be poorly edited. But is it so? What did actually shape selectivity of the editing domain? We used IleRS as a model enzyme and a series of amino acids of distinct physicochemical features and evolutionary origin to address these questions. We surprisingly found that tRNAs misaminoacylated with non-cognate amino acids that are efficiently discriminated at the activation step (**Table 1**) were all edited with the same rates as the biological threats Nva- and Val-tRNA^Ile^ (**Figure 2**). This demonstrates that recognition at the editing site is not influenced by how well the non-cognate amino acid is discriminated at the synthetic site. The similar was found for ValRS (6) and LeuRS (5, 22). It appears that the editing site is non-selective (except for the cognate AA-tRNA) and hydrolyses tRNAs misaminoacylated with amino acids spanning a broad range of physicochemical properties. How is this possible? The substrate recognition and catalysis at the class I editing site relies on the common parts of all AA-tRNAs; the terminal adenosine of the tRNA (A76) and α-NH_3_^+^ group of the amino acid attached to the tRNA anchor the substrates (42) while the 2’OH or 3’OH group of the A76 acts as a general base and promotes catalysis (shown for class I (16, 27) and class II (50, 51)). Changes of the terminal adenosine (52), lack of 2’OH or 3’OH (16, 27, 50, 51) or loss of the α-NH3+ anchoring interactions deprived editing (22, 53). Thus, it is plausible to assume that a common tRNA carrier and preselection of the editing substrates by the aminoacylation sieve, could have driven the evolution of the editing domain towards a broad substrate acceptance and the lack of recognition of the non-cognate amino acid’s side chain. In contrast, the synthetic site, acting as the first sieve, recognizes standalone amino acid and uses most of its side chain (**Table 1**) to minimize the error and ATP consumption (editing) (38). Thus, the synthetic and editing sites act as mirror images; while the former is highly selective to prevent errors, the latter exhibits low selectivity to clear each non-cognate amino acid that comes loaded to the tRNA.

Finding that class II PheRS, which recognizes the functional group of Tyr at the editing site (54) edits Ile-tRNA^Phe^ (55), indicates that broad selectivity is not confined only to the class I editing domain. Further, d-aminoacyl-tRNA deacylase, which bears a structural resemblance to the archaeal class II threonyl-tRNA synthetase editing domain (56), edits all d-amino acids at similar rates while efficiently rejecting l-amino acids. (57). In contrast, the editing domain (INS) of class II prolyl-tRNA synthetase (ProRS), as well as the free-standing bacterial ProRS INS domain homologs, have well-defined non-cognate amino acid specificity (58). Interestingly, the free-standing *trans*-editing proteins also evolved a broad specificity – in this case regarding the tRNA substrate (59). Thus, across the editing systems, a similar concept emerged independently arguing for the benefits of broad substrates acceptance in the design of the efficient error correction mechanisms.

### Negative catalysis ensures high specificity and broad selectivity of class I editing domain

Enzymes significantly differ in their physiological requirements for high selectivity (38). In some cases, low selectivity is beneficial allowing a broad substrate scope as in cytochrome P450 (60). The same applies to the editing domain. Yet, a unique feature of the editing domain, in which it mirrors highly selective enzymes (61), is its exquisite specificity *in rejection* of the cognate AA-tRNA. In general, specificity may evolve by positive and negative selection (21). While the former is a consequence of a selection for the enzyme’s high catalytic efficiency towards the cognate substrate, the latter is an explicitly evolved trait against a particular non-cognate substrate to avoid deleterious errors. Here we propose that specificity of the editing domain evolved through negative selection against the cognate AA-tRNA. We and others have previously shown that the cognate AA-tRNA is rejected from the editing site not by mitigating the binding, but by diminishing the catalysis (14, 16–18). Destabilisation of the transition state solely for cognate AA-tRNA hydrolysis can be viewed as an example of negative catalysis (20). This concept was introduced to explain that alongside promoting a wanted reaction by lowering the energy of the transition state for the desired product (positive catalysis), enzymes may also increase the energy barrier of the competing transition state preventing the side reaction (negative catalysis). Herein, we broaden this concept to compare transition states for the competing substrates. Thus, the residue conferring negative catalysis should not influence the rate of native (wanted) reaction but should diminish the reaction rate with the prohibited substrate. Visualisation of our data by activity-specificity graph revealed that the His333 and Thr246 IleRS residues confer negative catalysis (**Figure 6**). Their substitutions do not influence editing of Val-tRNAIle (*K*_Mut_/*k*_WT_ for editing is close to one) but promote misediting of Ile-tRNA^Ile^ resulting in the variants with decreased specificity (drop in the ratio of the *K*_Mut_/*k*_WT_ values for editing over misediting). In contrast, D342A mutation in IleRS promoted a decrease in both activity and specificity conferring the positive role for the Asp342 residue in catalysis (promoting both wanted and unwanted reaction by anchoring α-NH3 of the cognate and non-cognate amino acid (27, 42)). Similarly, LeuRS Thr252, which imposes unproductive positioning of Leu-tRNA^Leu^ (16), confers negative catalysis while the Asp345 residue (22), analogous to IleRS Asp342, promotes positive catalysis. In conclusion, negative selection/catalysis appears as a powerful mechanism to ensure low selectivity of the editing domain while keeping, at the same time, exquisite specificity in the rejection of the cognate AA-tRNA. The former was driven by relying on the common parts of all AA-tRNAs and the latter by the evolution of a specific kinetic rejection mechanism based on the cognate amino acid side chain (16–18).

**Figure 6.**
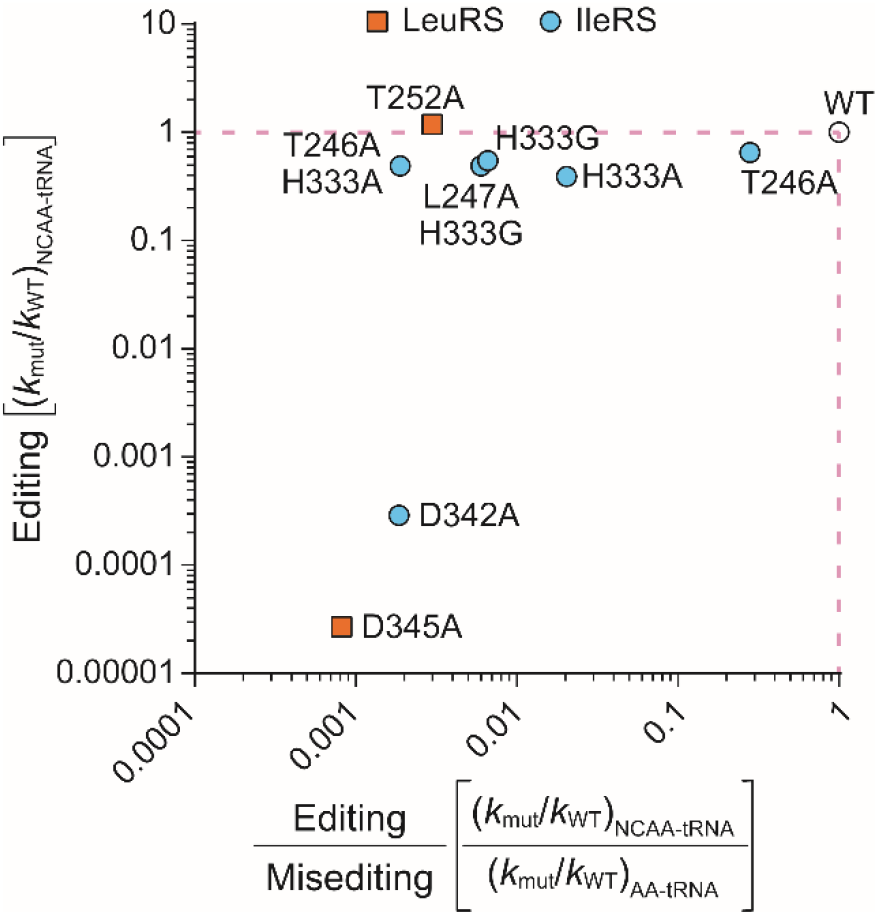
Activity-specificity relationship for editing of IleRS and LeuRS mutants. Activity (editing) was calculated as the ratio of the rate constants for hydrolysis of misaminoacylated tRNAs (NCAA-tRNA) by mutant AARS and WT, respectively. Specificity was calculated as the ratio of editing (wanted) over misediting (unwanted), where the misediting is the ratio of the rate constants for hydrolysis of cognate AA-tRNAs by mutant AARS and WT, respectively.

### IleRS – class Ia enzyme with unique editing features

In class I AARSs, delivery of the amino acid from the synthetic to the editing site occurs through fast translocation (estimated as faster than 80 s^−1^ in LeuRS (22)) of the 3’-end of the (mis)aminoacylated tRNA (62) (**Figure 1**). The fast translocation allows that the 3’-end reaches the editing site on a shorter time scale relative to the rate-limiting dissociation of the (mis)aminoacylated tRNA. Therefore, editing occurs *in cis* without the release of misaminoacylated tRNA. Release and rebinding of the misaminoacylated tRNAs to AARS was shown so far only for class II AARS (14), and was attributed to fast dissociation of (mis)aminoacylated tRNA (3). It, therefore, came as a surprise that IleRS edits Val-tRNA^Ile^ with a significant contribution of the *in trans* pathway. At the same time, both LeuRS and ValRS predominantly operate *in cis* (**Figure 5**). Thus, IleRS appears unique among closely related class Ia editing AARSs (ILVRS) in post-transfer editing. Although the mechanistic basis is still missing, it is worth commenting here that only IleRS, among ILVRS, uses tRNA to both optimize the affinity for the amino acid substrate and to stimulate pre-transfer editing (**Figure 1, path 3**). Indeed, *E. coli* IleRS showed substantial tRNA-dependent pre-transfer editing that comprises about 30 % of total editing (11, 22, 37). Thus, an idiosyncratic utilisation of the tRNA as a co-factor may influence the rates of the tRNA-dependent steps, for example translocation.

It has been shown that PheRS can compete with EF-Tu for binding to Tyr-tRNA^Phe^ (14) providing a basis for editing *in trans*. In our hands, EF-Tu efficiently competes with IleRS for binding to Val-tRNA^Ile^, but the experimental set up was designed to promote the EF-Tu interaction. Further, this *in vitro* observation is distinct from *in vivo* environment where other AA-tRNAs compete for EF-Tu. Nevertheless, it is plausible to assume that editing *in trans* is generally of lower proficiency than editing *in cis*, because EF-Tu may bind tRNAs misaminoacylated with amino acids similar to the cognate one (14, 23, 44) and redirect them to ribosomal translation. That said, IleRS capacity to edit errors prior misaminoacylation could be relevant opening a provocative question to what extent is AA-tRNA leakage and its editing *in trans* intertwined with pre-transfer editing.

## AVAILABILITY

All data are available from the authors upon request.

## SUPPLEMENTARY DATA

Supplementary Data are available at NAR online.

## ACKNOWLEDGEMENT

We gratefully acknowledge Marko Mocibob, Vladimir Zanki and Vijay Jayaraman for fruitful discussions and comments regarding this manuscript. Ita Gruic-Sovulj dedicates this manuscript to the memory of Dan Salah Tawfik, whose generosity, creativity, and passion for life and science will continue to inspire many of us.

