## Supplementary material for "Negative catalysis by the editing domain of class I aminoacyl-tRNA synthetases": SI Appendix

### Supplementary data

#### Figures

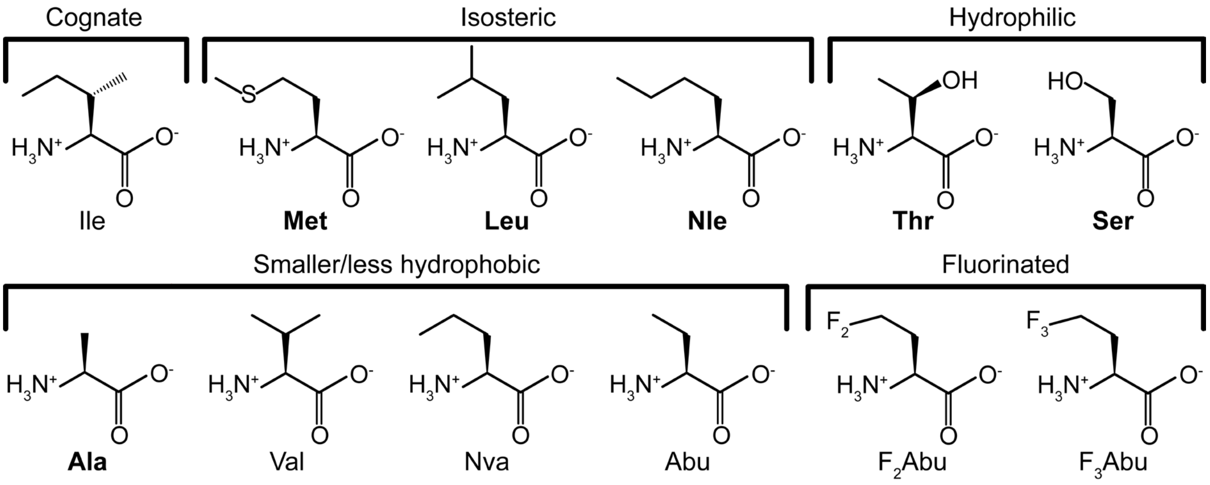


Figure S1. Structures of the amino acids tested as non-cognate substrates of IleRS grouped by their physicochemical properties: methionine (Met), leucine (Leu), norleucine (Nle), threonine (Thr), serine (Ser), alanine (Ala), valine (Val), norvaline (Nva), α-aminobutyrate (Abu), α-amino-γ,γ-difluorobutyrate ($F_2$Abu), and α-amino-γ,γ,γ-trifluorobutyrate ($F_3$Abu). Isoleucine (Ile) is the cognate substrate. The bolded amino acids were analysed in this work. We assign Met, Leu and Nle as isosteric to Ile as they share the same volume (Alvarez-Carreño et al., 2013).

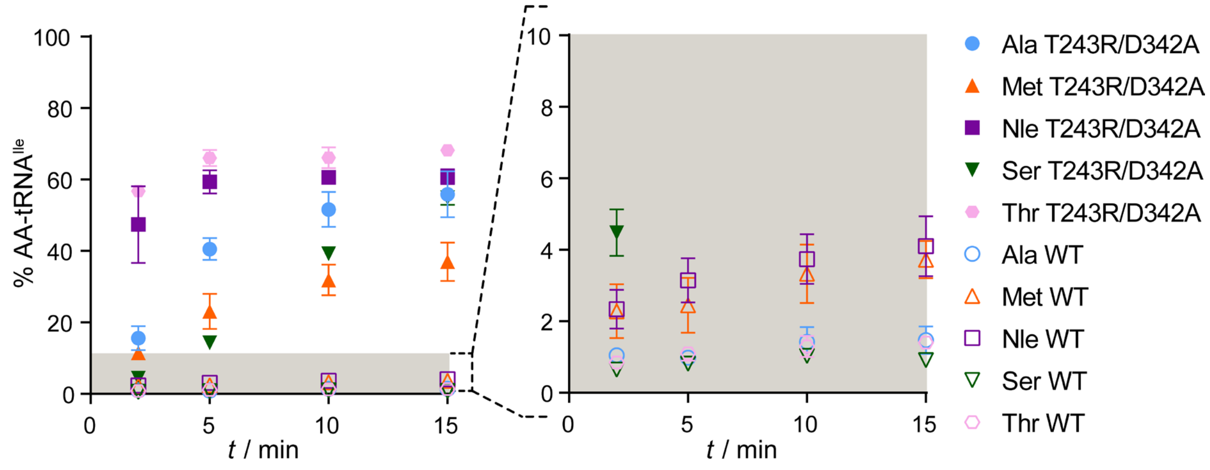


Figure S2. Misaminoacylation of $tRNA^{Ile}$ with different non-cognate amino acids by 1 µM post-transfer editing deficient T243A/D342A IleRS and WT IleRS (inset). Amino acids were present at a moderate amount to minimise aminoacylation of Ile, which can be potentially present in trace amounts in the non-cognate amino acid samples (Nle and Thr 2 mM; Ala and Met 4 mM; Ser 10 mM).

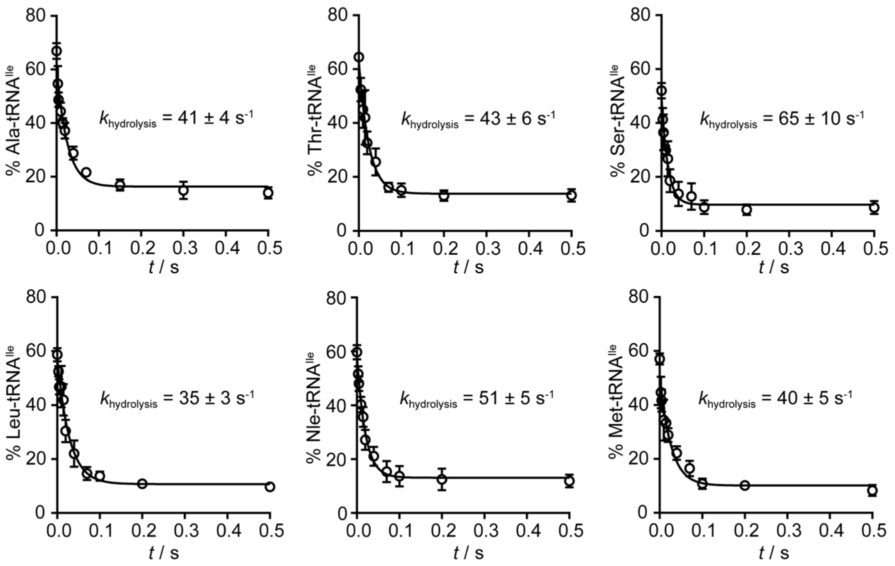


Figure S3. Single-turnover time courses for hydrolysis of misaminoacylated $tRNA^{Ile}$s by WT IleRS. The time course data fit the best to monoexponential decay allowing calculation of $k_{hydrolysis}$ – the first-order rate constant for AA-tRNA hydrolysis within the editing site. Time points represent mean value ± SEM for three independent experiments. High Y values at $t$ = 0 demonstrates the efficiency of preparative tRNA misaminoacylation using post-transfer editing deficient T243R/D3432A IleRS (please note high levels of Met-, Leu- and Nle-$tRNA^{Ile}$ despite possible Ile contaminations of these amino acids samples).

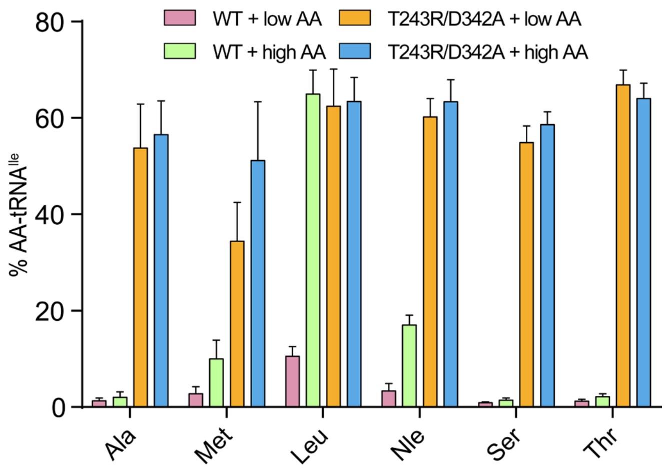


Figure S4. Assessment of the purity of the amino acid samples. The purity was tested by misaminoacylation of 5 µM $tRNA^{Ile}$ with two different concentrations of the non-cognate amino acids (Leu 0.2 and 2 mM; Nle and Thr 2 and 20 mM; Ala and Met 4 and 40 mM, Ser 10 and 100 mM) using WT and post-transfer editing deficient T243R/D342A IleRS. The Y-axis values represent the plateau of AA-$tRNA^{Ile}$ achieved after 15 minutes of aminoacylation. WT enzyme accumulates AA-tRNA in the presence of Met, Nle and Leu, which may originate from aminoacylation of Ile traces present in the amino acid samples (Ile-$tRNA^{Ile}$ is hydrolysed by two orders of magnitude slower rate than other misaminoacylated tRNAs, Figure 2). In the case of Leu, the plateau of AA-tRNA increases 5-fold with a 10-fold increase in the concentration of Leu, which indicates contamination. In the case of Met and Nle, the situation is more ambiguous, but the contamination cannot be completely excluded. The contamination could compromise the calculation of $D$, resulting in an artificially lower level of discrimination and therefore $D$ was not calculated (Table 1).

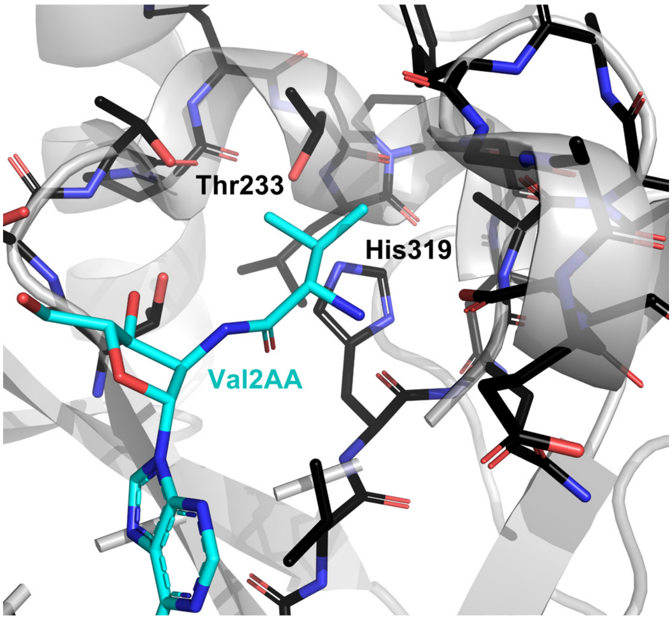


Figure S5. Structure of the *T. thermophilus* IleRS editing site with the post-transfer editing reaction analogue 2'-(L-valyl)amino-2'-deoxyadenosine bound (PDB ID: 1WNZ).

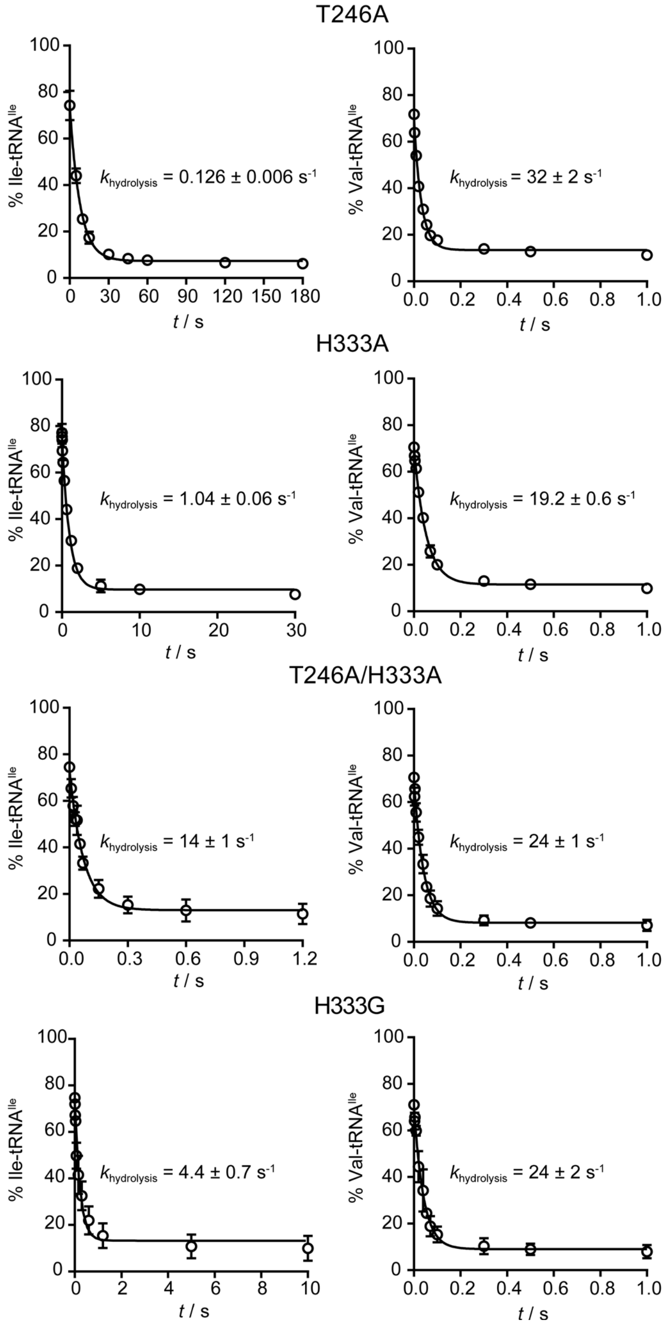


Figure S6. Single-turnover time-courses for hydrolysis of Ile- and Val-$tRNA^{Ile}$ by T246A, H333A, T246A/H333A and H333G IleRS. The time course data fit the best to monoexponential decay allowing calculation of $k_{hydrolysis}$ – the first-order rate constant for AA-tRNA hydrolysis within the editing site. Time points represent mean value ± SEM for three independent experiments.

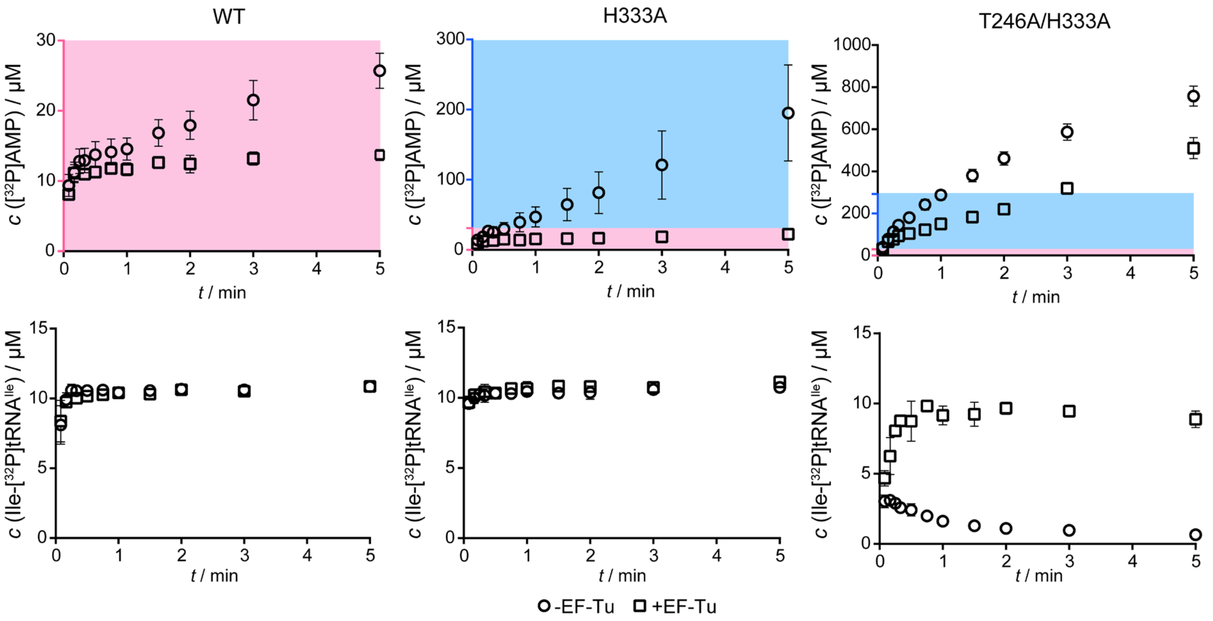


Figure S7. ATP consumption (represented by AMP formation at upper panels) and Ile-tRNA$^{Ile}$ formation (lower panels) were followed in parallel reactions (the former using [$^{32}$P]ATP and the later [$^{32}$P]tRNA) with (circles) and without (squares) 8-12 µM active EF-Tu. IleRS and its variants were present at 2 µM, tRNA$^{Ile}$ was 12 µM. Background colours indicate the difference in the y-scale.

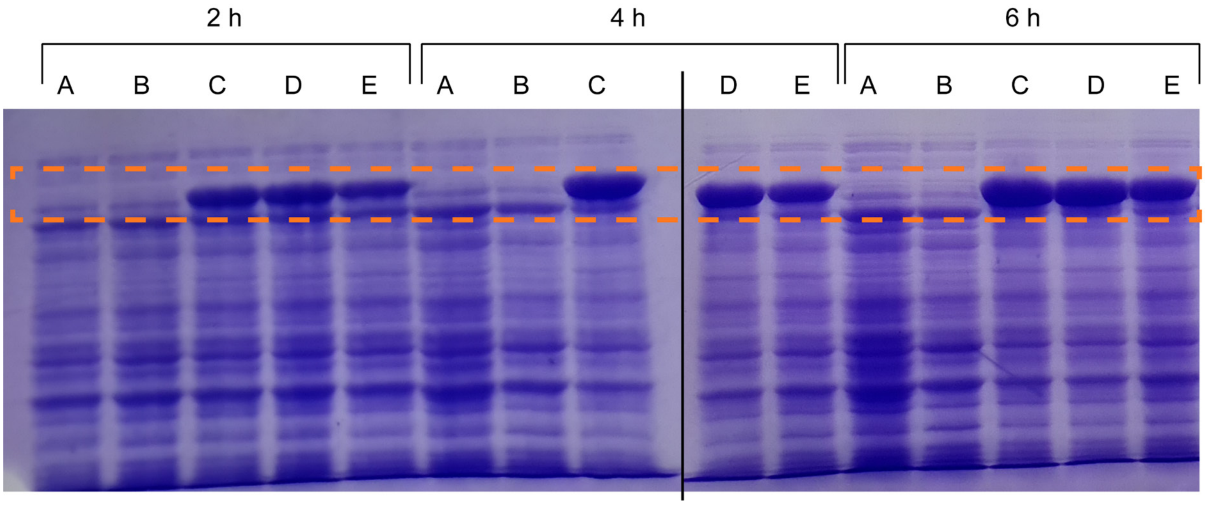


Figure S8. Expression of IleRS variants during the growth of *E. coli* Bl21 transformed with pET28b carrying *ileS* variants. A – pET28b, B – pET28b plus 100 µM IPTG, C – pET28b_*IleS* plus 100 µM IPTG, D – IleRS H333A and E – IleRS T246A/H333A. IleRS band is denoted by the dashed orange rectangle. The black vertical line separates two SDS-PAGE gels. Each well contained the same amount of cells (equivalent to 1 ml of cell suspension at $OD_{600}$ = 0.1.)

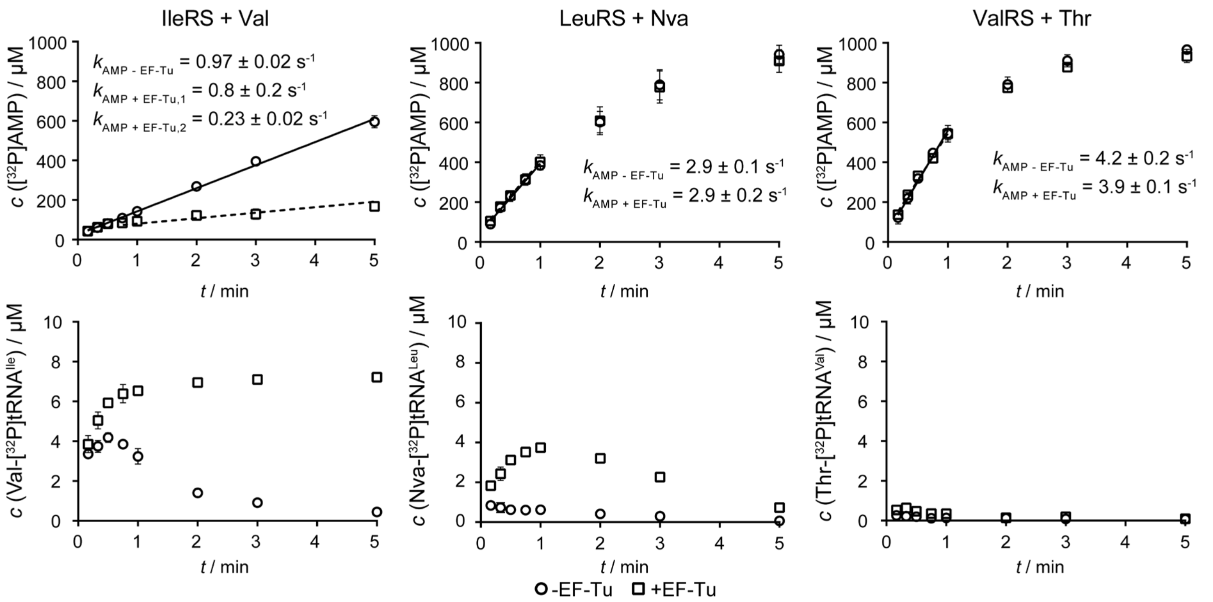


Figure S9. ATP consumption (represented by AMP formation at upper panels) and AA-tRNA formation (lower panels) were followed in parallel reactions (the former using [$^{32}$P]ATP and the later [$^{32}$P]tRNA) with (circles) and without (squares) 8-12 µM active EF-Tu. AARS were present at 2 µM, tRNA$^{Ile}$ was 12 µM.

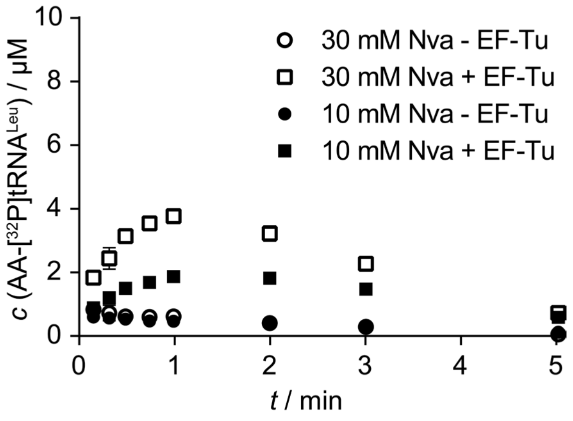


Figure S10. Aminoacylation of the tRNA$^{Leu}$ in the presence of 10 and 30 mM Nva by WT LeuRS with and without 8-12 active EF-Tu. AARS were present at 2 µM, tRNA$^{Ile}$ was 12 µM. The aminoacylation level depends on the Nva concentration suggesting that Leu-tRNA$^{Leu}$ is mainly followed (Leu comes from trace contaminations of Nva). The Nva-tRNA$^{Leu}$ aminoacylation level should not strongly depend on the increase in Nva concentration due to saturated conditions.
